# Adolescent morphine exposure does not alter low-dose lipopolysaccharide-induced sickness behavior in adult mice

**DOI:** 10.1101/2025.06.26.661880

**Authors:** Natalie V. Davidson, Cynthia Masese, Caitlin Han, Lili Massac, Yuu Ishikawa, Grace Reynolds, Codey Smithson, Jackson Cook, Melissa T. Manners, Shivon A. Robinson

## Abstract

Adolescent opioid use in the United States commands attention: millions of twelve- to nineteen-year-olds are exposed to opioids each year by prescription and misuse. Recent findings show that opioids bind not only to canonical opioid receptors but also interact with receptors on immune cells within both the central and peripheral nervous systems. The potential for early life opioid exposure to induce long-term changes in the neuroimmune system is not fully understood, particularly given the adolescent brain’s high susceptibility to neuroplastic changes. The goal of this study was to investigate the hypothesis that adolescent opioid use potentiates physiological and behavioral responses to pathogen-induced sickness later in life. To achieve this, we treated adolescent mice (PND 35-42) with bi-daily escalating doses of morphine to model dependence and then administered a low dose of lipopolysaccharide (LPS, 0.1 mg/kg) in adulthood (PND 60-76) to induce an immune response. In contrast to our hypotheses, we found that adolescent morphine exposure had no additive effect on low-dose LPS-induced sickness measures when assessed in adulthood. These data suggest that adolescent opioid exposure may have minimal effects on future immune challenges, although further research is needed to confirm this.

## Introduction

Despite the declining rate of dispensed opioid prescriptions (Zhang 2023) and opioid-related overdoses (Ahmad 2025), opioid misuse and addiction remain critical public health issues in the United States. A growing concern within the current opioid crisis is the rising cases of adolescent opioid use. In 2023, approximately 5.7 million Americans aged 12 and older had an opioid use disorder, and 8.6 million reported misusing prescription painkillers [1]. Notably, a multi-center retrospective study reported that among 12-17 year-olds, the incidence of opioid-related presentations to the emergency department increased 2800% between 2014 and 2022 [2]. Nonmedical use of prescription opioids during adolescence is correlated with high-risk behaviors, such as violence victimization, poor academic performance, and suicidal thoughts [3]. Furthermore, adolescent opioid use is associated with increased risk of developing opioid dependence and substance use disorder symptoms later in life [4,5]. Because adolescence is a critical neurodevelopmental period in which individuals may be more susceptible to the effects of psychoactive substances on brain functioning, [6–8] there is a pressing need to further elucidate the neurobiological impact of adolescent opioid exposure on long-term health outcomes.

Opioids classically exert their analgesic and euphoric effects via binding to endogenous opioid receptors located throughout the central and peripheral nervous system [9–11]. Opioids can also modulate immune signaling via interaction with opioid receptors expressed on various immune cells, including lymphocytes and macrophages [12–14]. Previous studies have demonstrated that opioids have immunosuppressive effects in the periphery [15–18]. However, there is evidence that opioids can bind to the accessory proteins of the Toll-Like Receptor 4 (TLR4) [19,20], which is canonically activated by pathogen-associated endotoxins, such as lipopolysaccharide (LPS), and stimulates the production of proinflammatory cytokines [21,22]. Opioid-mediated activation of TLR4 receptors within the central nervous system (CNS) has been shown to induce neuroinflammation [20,23,24].

Elevated pro-inflammatory cytokine signaling in the CNS triggers complex neuroimmune mechanisms that manifest as distinct physiological and behavioral responses, often referred to as “sickness behaviors” [25,26]. In rodents, LPS-induced sickness is characterized by decreased motor activity, weight loss, body temperature changes, and depressive-like behaviors [27–30], though many of these effects are dose- and time-dependent. Sickness behaviors can be adaptive and promote recovery by shifting energy allocation toward fighting infection [31]. However, prolonged or excessive immune activation may have detrimental outcomes. For example, chronic neuroinflammation is associated with the development of depression [32–34] and has been proposed as a key factor in modulating the impacts of social-environmental adversity and stress on overall health and lifespan [35].

Previous studies have demonstrated that chronic opioid exposure dysregulates immune system functioning, potentially leading to increased susceptibility to infection and maladaptive immune responses [12,36–38]. Notably, administration of a mu-opioid receptor antagonist reduces LPS-induced inflammation and sickness behavior in mice [39–41]. While the majority of this work has been conducted in adult models, there is growing evidence indicating that early-life opioid exposure is disruptive to future immune functioning. In humans, prenatal opioid exposure is associated with increased risk of childhood infection [42]. In rodents, perinatal morphine exposure has been shown to exacerbate LPS-induced neuroinflammatory responses and sickness behaviors in adulthood [43,44].

Adolescence is also a vulnerable developmental period for immune insults [45–47]. It has been demonstrated in rats that adolescent morphine exposure potentiates morphine-induced neuroimmune activation and drug-seeking behavior in adulthood [48]. However, there is limited preclinical investigation into whether adolescent opioid exposure specifically alters LPS-induced sickness behaviors later in life. To investigate this, we administered morphine to male and female adolescent mice using a 5-day escalating dose paradigm. In adulthood, the mice were exposed to a low dose of LPS (0.1 mg/kg) and assessed for physiological and behavioral signs of sickness, including alterations in body weight, body temperature, locomotion, and behavioral despair as measured by the forced swim test (FST). We hypothesized that adult animals with a history of adolescent morphine exposure would exhibit more severe LPS-induced sickness symptoms compared to controls.

## Materials and Methods

### Animals

Male and female C57BL/6NTac mice (n = 66) obtained from Taconic Biosciences were used for all experiments. Mice were maintained on a 12-hour dark-light cycle (lights on at 07:00) in a temperature and humidity-controlled facility with food and water available ad libitum. All experiments were performed in compliance with the National Institutes of Health’s Guide for Care and Use of Laboratory Animals, and protocols were approved by the Institutional Animal Care and Use Committee at Williams College.

### Drug treatment

Morphine sulfate was obtained from Millipore Sigma (St. Louis, MO) and dissolved in 0.9% sterile saline. Adolescent mice (PND 35-42) received repeated subcutaneous injections of morphine beginning on the evening of Day 1 and then twice daily (at 9 AM and 4 PM) at escalating doses (10, 20, 30, 40, or 50 mg/kg), with the final injection occurring on the morning of Day 5. An escalating dosing schedule has previously been shown to induce dependence in adolescent animals [49]. Controls received saline at identical timepoints. Animals were weighed daily during morphine treatment and up to 1 week after the final injection. Following adolescent morphine exposure, mice were left undisturbed until adulthood.

Lipopolysaccharide (LPS) derived from *E. coli* 0111: B4 was obtained from Millipore Sigma (St. Louis, MO) and dissolved in 0.9% sterile saline. During adulthood (PND 60-67), mice were administered a single intraperitoneal injection of LPS (0.1 mg/kg) or saline. This dose has previously been shown to elevate pro-inflammatory cytokine levels and induce mild to moderate sickness symptoms [30,50] while having minimal effects on affective behaviors in the absence of an additional insult [51]. Following LPS administration, mice were assessed for several physiological and behavioral signs of sickness. An overview of the experimental timeline is shown in Fig. 1.

**Figure 1.**
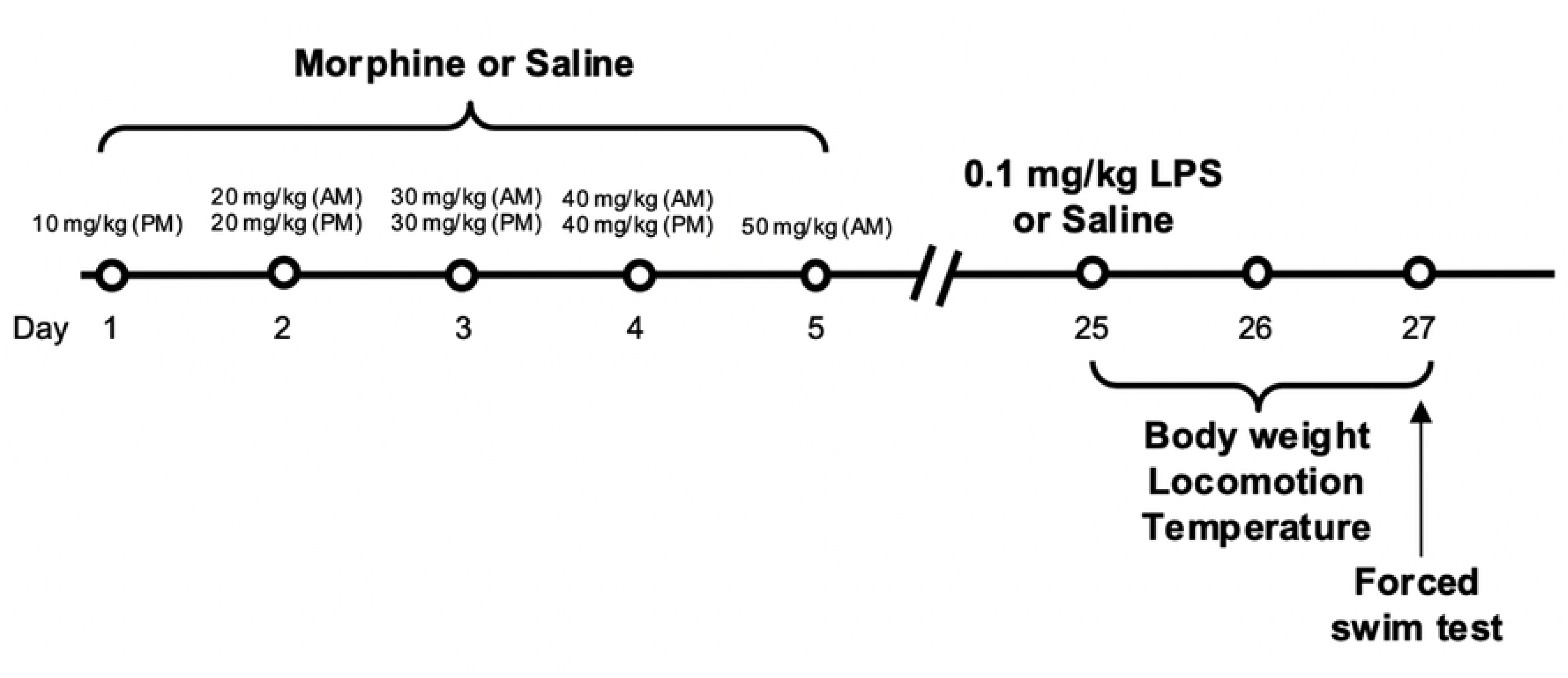
Experimental timeline.

All drugs were delivered in a 10 ml/kg injection volume, and controls received sterile saline in an equal injection volume.

### Physiological measures

Animals were weighed immediately before the LPS or saline injection, and then 24 and 48 hours following the injection. Body surface temperature was measured via an infrared thermometer (iProven) aimed at the lower abdomen [52] immediately before LPS or saline administration, and then 1, 24, and 48 hours after administration. Body weight and surface temperature were calculated as percent change from baseline.

### Locomotor activity

The effect of LPS on locomotor activity was assessed 1, 24, and 48 hours after administration. Mice were habituated to the experimental room for 30 minutes before testing and then individually placed in a 20 x 20 x 35 cm acrylic apparatus for 30 minutes. Distance traveled in meters was measured using ANY-maze video tracking software (Stoelting Co., Wood Dale, IL).

### Forced swim test

Behavioral despair was assessed utilizing the forced swim test (FST) 52 hours after LPS administration. This time point was selected based on preliminary data indicating that the locomotor suppressing effects of LPS dissipated by 48 hours post-treatment. Mice were habituated to the experimental room for 30 minutes before testing and then placed individually in a plastic cylinder (26cm tall x 18 cm diameter) filled with water (24 °C +/- 1 °C) to a depth of 15 cm for six minutes. Total time immobile in the last 4 minutes of the 6-minute test was measured using Anymaze tracking software (Stoelting Co., Wood Dale, IL). A period of immobility was defined as 2 seconds or greater of no movement [53].

### Statistical analysis

Data were analyzed and graphed using the package “afex” within R and GraphPad Prism 9 software, respectively. Data were analyzed using linear models including sex, morphine treatment, LPS treatment, and the interaction term as fixed effects. Linear mixed-effects models were used for datasets that included repeated measurements, and also included timepoint (Day) as a fixed effect and subject as a random effect. Statistical outliers were identified using the two-sided Grubbs’ test (α = 0.05) and excluded from analysis. Post-hoc analyses were performed using Bonferroni’s post hoc tests. Statistical significance was set at p < 0.05 throughout.

## Results

### Adolescent morphine exposure reduces body weight

Adolescent male and female mice were administered bi-daily escalating doses of morphine over five days to induce morphine dependence. We observed a significant main effect of morphine treatment [F(1, 61) = 116.83, p < 0.0001] and time [F(5, 305) = 174.65 p < 0.0001], in addition to a significant morphine treatment * time interaction [F(5, 305) = 53.96, p < 0.0001] on body weight. Post-hoc analyses revealed that morphine-treated animals exhibited greater weight loss than saline-treated animals from Day 2 to Day 12, indicating that morphine’s effect on body weight persists up to 1 week after treatment (Fig. 2).

**Figure 2.**
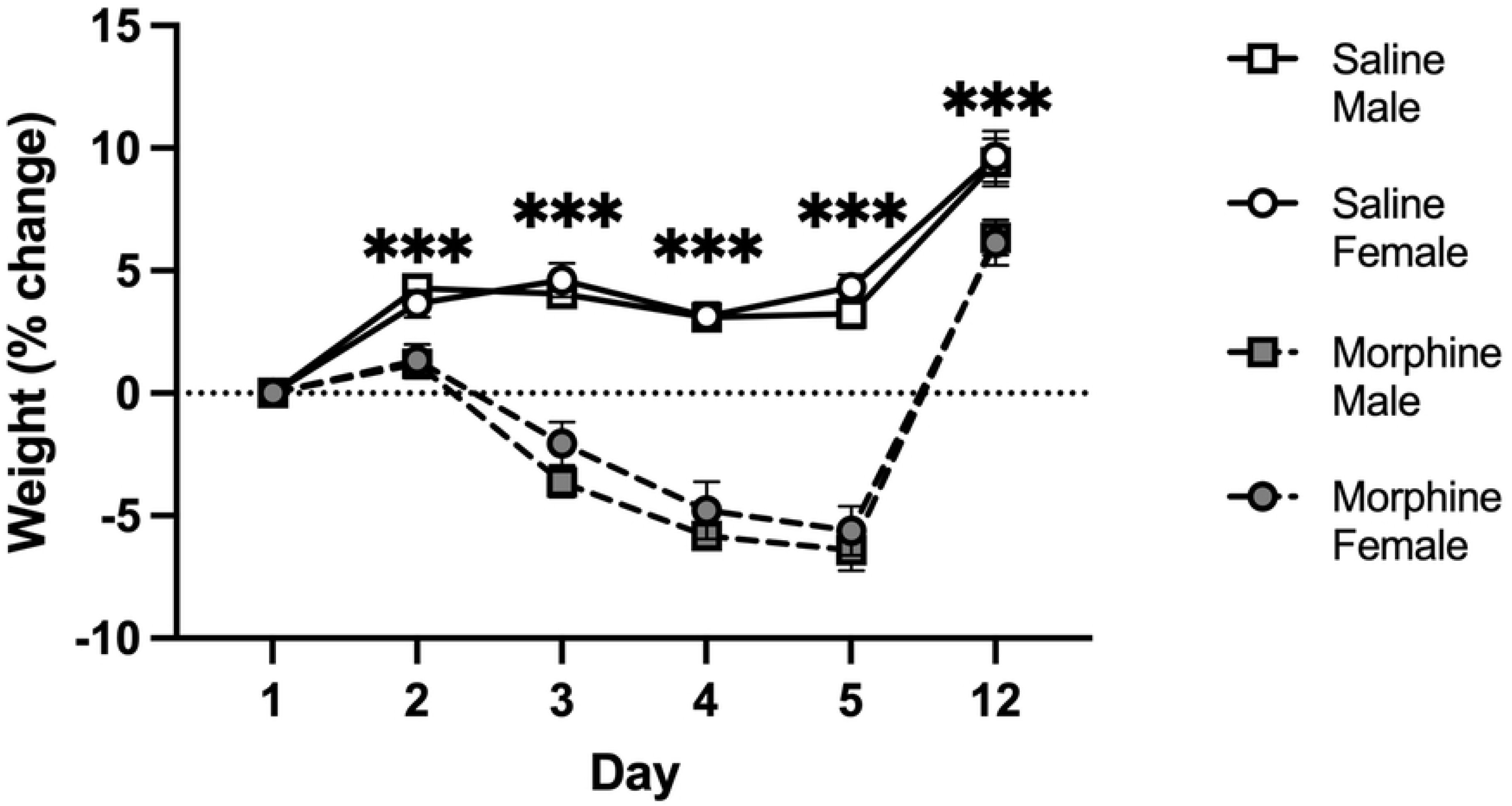
Adolescent morphine exposure blunts weight gain. Adolescent mice received bi-daily escalating doses of morphine or saline for 5 days. Body weight was measured each day of drug treatment (Days 1-5) and one week after the final drug administration (Day 12). Morphine-treated mice exhibited greater weight loss than saline-treated mice between Day 2 and Day 12 of treatment. Saline, *n* = 18 males, 16 females; morphine, *n* =15 males, 16 females. Data are presented as mean ± SEM. ***p < 0.001 compared to saline-treated.

### LPS treatment reduces body weight and surface temperature, independent of morphine exposure

There was a significant main effect of LPS treatment [F(1,55.97) = 75.33, p < 0.0001], time [F(1, 54.30) = 115.40, p < 0.0001], and a LPS treatment * time interaction [F(1,54.30) = 138.80, p < 0.0001] on body weight. Post-hoc analyses revealed that LPS-treated animals weighed less than saline-treated animals 24 hrs (p < 0.0001) and 48 hrs (p < 0.001) post-treatment (Fig. 3a). Furthermore, we observed a significant morphine treatment * LPS treatment interaction [F(1,55.97) = 4.19, p < 0.05], sex * time interaction [F(1,54.30) = 7.10, p < 0.05], and a sex * LPS * time interaction [F(1,54.30) = 7.24, p < 0.01]. However, no significant differences were detected following post-hoc analyses.

**Figure 3.**
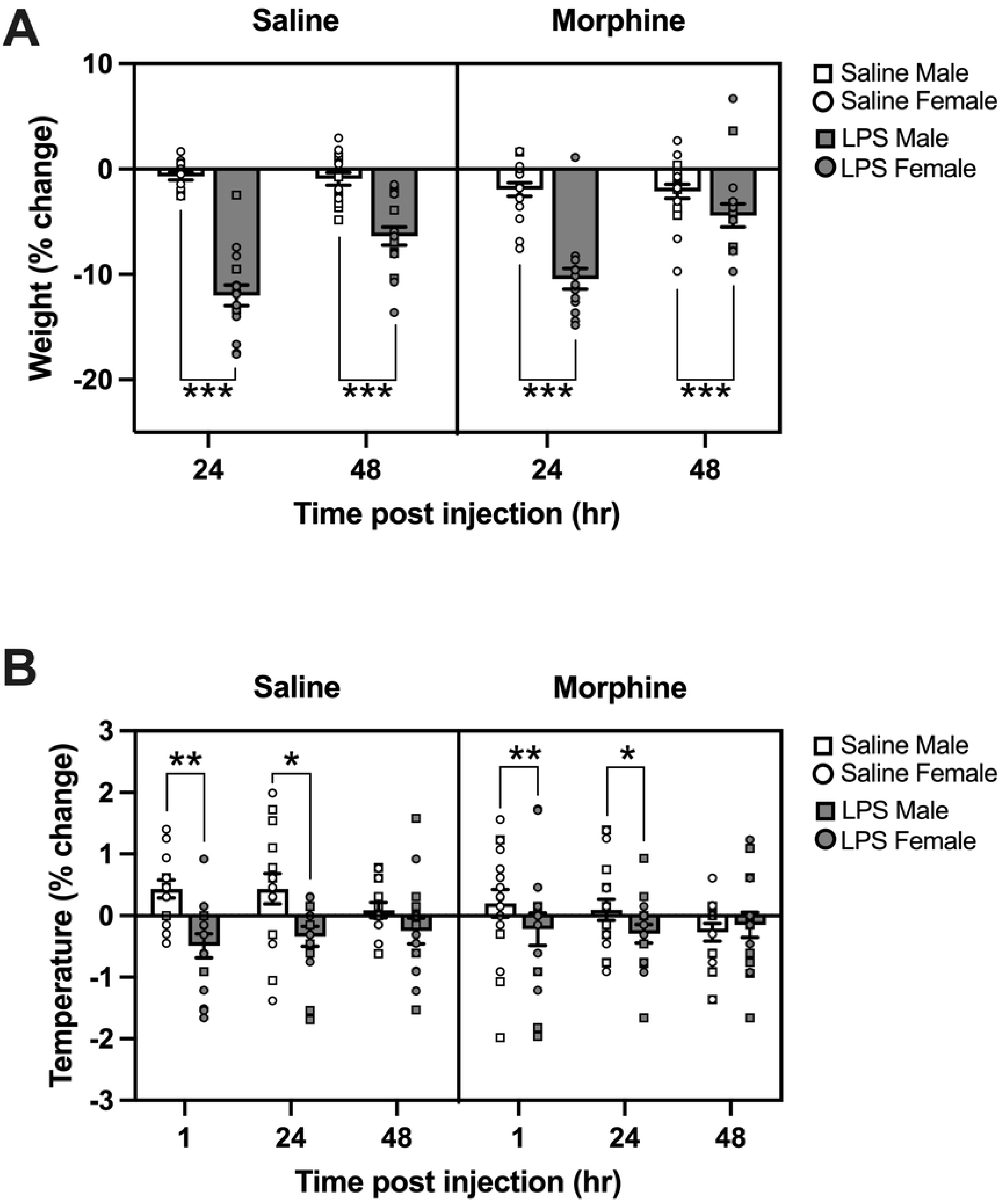
LPS administration reduces body weight and temperature. Mice previously exposed to either saline or morphine during adolescence were later administered LPS (0.1 mg/kg) or saline in adulthood. A) Baseline body weight was recorded and then measured 24- and 48-hours post-injection. Mice treated with LPS exhibited significantly more weight loss than saline-treated animals at 24 and 48 hours. B) Abdominal surface temperatures were measured at baseline, then 1, 24, and 48 hrs post-injection. LPS-treated mice exhibited significantly reduced abdominal surface temperature compared to saline-treated mice 1 hour and 24 hours post-injection. Saline-saline, *n* = 7 males, 8 females; saline-LPS, *n* = 8 males, 8 females; morphine-saline, *n* = 10 males, 8 females; morphine-LPS, *n* = 8 males, 8 females. Data are presented as mean ± SEM. ***p < 0.001, *p < 0.05 compared to saline-treated.

Surface temperature was measured 1, 24, and 48 hours post-LPS injection by aiming an infrared thermometer at the abdomen. There was a significant main effect of LPS treatment [F(1,53.09) = 9.53, p < 0.01] and a LPS treatment * time interaction [F(2,106.93) = 3.88, p < 0.05] on surface temperature alterations. Post-hoc analyses revealed that body temperature was reduced in LPS-treated animals at 1 hr (p < 0.01) and 24 hrs (p < 0.05) after injection; however, this difference dissipated by 48 hours (Fig. 3b).

### LPS treatment reduces locomotor activity independent of morphine exposure

Mice were assessed for alterations in locomotor activity 1, 24, and 48 hours after LPS administration. Analyses revealed a significant main effect of LPS [F(1,56.53) = 25.06, p < 0.001], time [F(2,111.10) = 56.73, p < 0.0001], and LPS * time interaction [F(2,111.10) = 48.00, p < 0.0001]. Post-hoc analyses revealed that LPS treatment reduced locomotor activity 1 hr (p <0.0001) and 24 hours (p < 0.05) post-administration (Fig. 4). Additionally, there was a significant morphine * time interaction [F(2,111.10) = 3.27, p < 0.05]; however, no significant differences were detected in post-hoc analyses.

**Figure 4.**
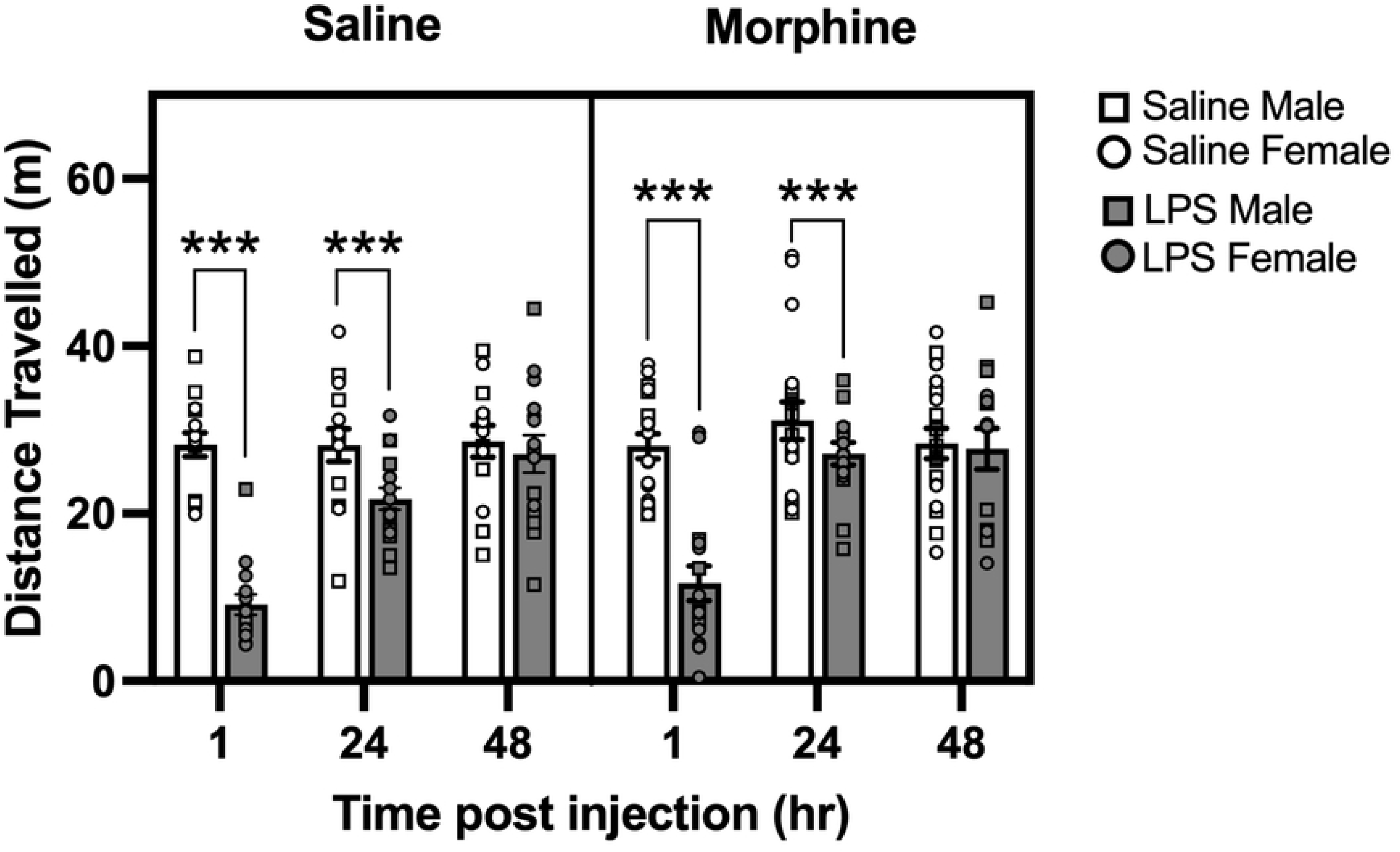
LPS administration reduces locomotor activity. Locomotor activity was assessed 1, 24, and 48 hrs after LPS (0.1 mg/kg) or saline administration. Mice treated with LPS travelled less than saline-treated animals when measured 1 hour and 24 hours post-injection. Saline-saline, *n* = 7 males, 8 females; saline-LPS, *n* = 8 males, 8 females; morphine-saline, *n* = 10 males, 8 females; morphine-LPS, *n* = 8 males, 8 females. Data are presented as mean ± SEM. ***p < 0.001 compared to saline-treated.

### No effect of drug treatment on behavior in the forced swim test

At 52 hours post-LPS injection, mice were assessed for time spent immobile in the forced swim test. No significant effects were observed for morphine treatment [F(1, 56) = 3.18, p = 0.08], LPS treatment [F(1 56) = 0.10, p = 0.75.] or sex [F(1, 56) = 0.02, p = 0.89] (Fig. 5).

**Figure 5.**
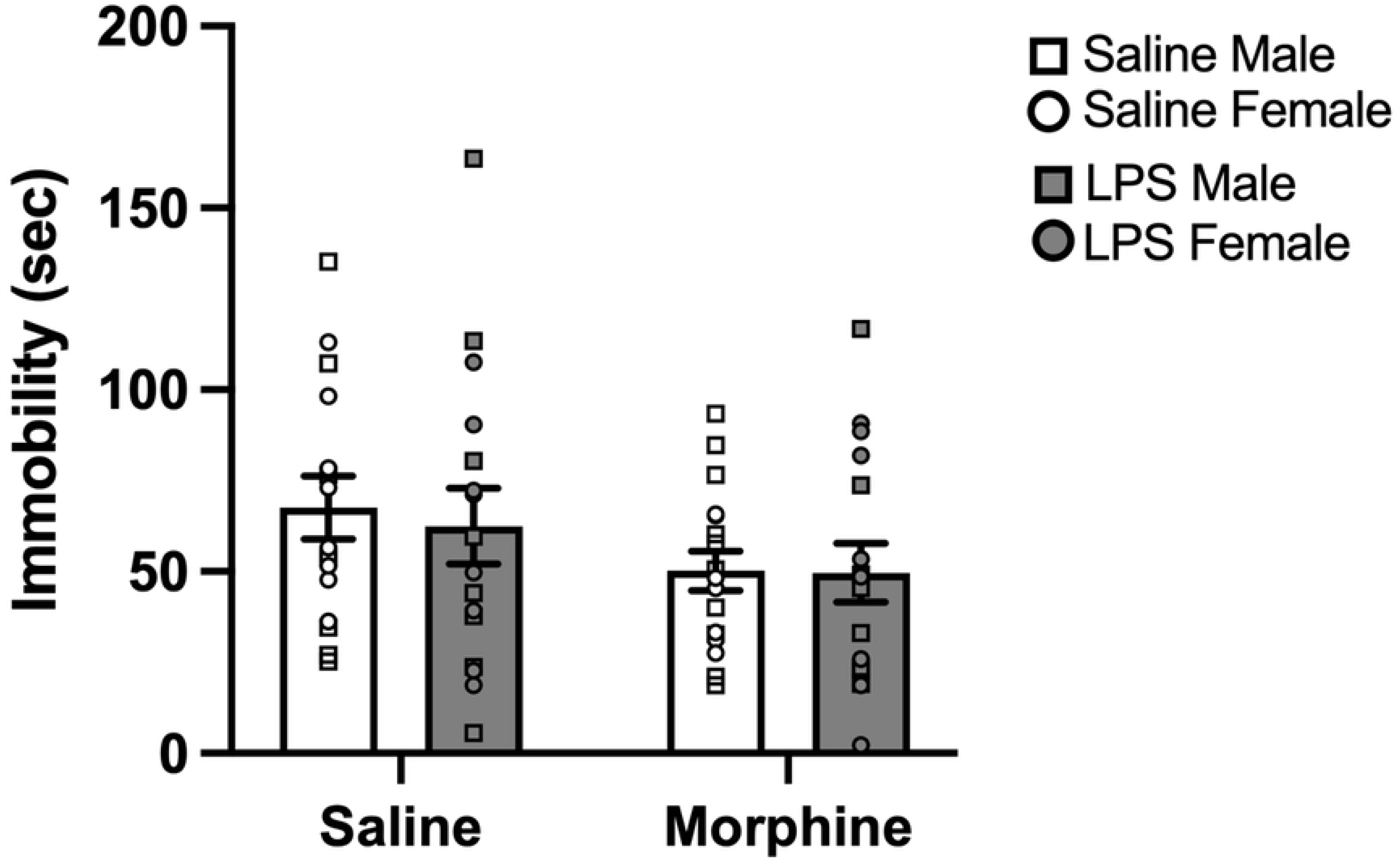
No effect of drug treatment on forced swim test. Mice were assessed in the forced swim test 52 hours post-LPS administration. There were no differences across treatment groups or sex in time spent immobile. Sal/Sal = 7M/8F, Sal/LPS = 8M/8F, Mor/Sal = 10M/8F, Mor/LPS = 8M/8F. Data are presented as mean ± SEM.

## Discussion

There is mounting clinical and preclinical evidence indicating that early-life opioid exposure can alter immune activation later in life [42–44,48]. The goal of the present study was to determine whether adolescent morphine exposure in mice alters sickness behaviors in response to an immune challenge in adulthood. Morphine dependence was achieved utilizing an escalating dosing schedule previously shown to produce robust autonomic and somatic withdrawal symptoms in adolescent mice [49]. Weight loss is a well-documented marker of morphine-induced metabolic changes in rodents [54,55], and adolescent morphine exposure has previously been shown to blunt weight gain in mice [56]. Consistent with these findings, we observed that mice in the adolescent morphine treatment group showed a greater reduction in weight compared to saline controls beginning on the second day of drug administration and lasting up to at least a week after the cessation of treatment. This suggests that the morphine dose and administration paradigm utilized in this study effectively induced physiological changes.

Lipopolysaccharide (LPS), an endotoxin found on the outer membrane of gram-negative bacteria, is a commonly used tool to investigate immune response in rodents due to its ability to activate pro-inflammatory cytokine signaling pathways and induce many symptoms of sickness observed in humans and animal models, including weight loss, lethargy, and alterations in mood, among others [26,27]. We predicted that morphine pre-exposed mice would exhibit a potentiated sickness response to a low-dose LPS (0.1 mg/kg) challenge. Contrary to our hypotheses, all LPS-treated animals exhibited similar symptoms of LPS-induced sickness regardless of adolescent drug exposure. LPS treatment significantly reduced body weight for up to 48 hours post-administration and suppressed locomotor activity for up to 24 hours post-administration. Interestingly, LPS treatment reduced body surface temperature up to 24 hours post-administration, in contrast to the febrile response often associated with systemic inflammation [57]. This could be due to surface-level vasoconstriction, a known response to LPS in mouse tail skin that coincides with a rise in core body temperature [58]. Alternatively, this finding may indicate a hypothermic core temperature response to LPS, which has previously been reported in mice [59,60] and may be a protective mechanism against LPS-induced shock [61].

In the present study, body temperature was measured using an infrared thermometer, which is considered to be a reliable non-invasive method for assessing body temperature in mice [52,59]. However, while potentially more distressing to animals, measuring core temperature via a rectal thermometer or implantable probe may have uncovered interactions between adolescent opioid exposure and adult sickness symptoms. Indeed, data collected from temperature sensors implanted in the abdominal cavity revealed that perinatal morphine exposure potentiated LPS-induced fever in adult female rats [44]. Furthermore, since implantable thermometers can collect more frequent data points over an extended period, this method may be better suited for detecting subtle or temporal effects.

Chronic neuroinflammation is associated with anxiety and depression-like symptoms in rodents [32–34]. A septic dose of LPS (5 mg/kg) has been shown to increase anxiety and behavioral despair in mice up to a month after treatment [62]. Notably, a low dose of LPS (0.1 mg/kg) combined with a chronic stress paradigm induces depressive like behavior in mice, whereas LPS treatment alone has no effect [51]. Despite these findings, we observed no differences in forced swim test (FST) behavior among treatment groups. This may be due to the timing of behavioral testing. We conducted the FST assay at 52 hours after establishing that LPS-treated animals no longer exhibited suppressed locomotor activity. However, potential depressive-like effects of LPS may have dissipated by this time point. Previous studies have reported LPS-induced impact on anxiety and depressive-like behavior as early as 24 hours post-administration [51,53]. More work is needed to fully characterize the temporal profile of LPS-induced affective behavior in this model.

Recent studies have reported changes in neuroimmune-related adult behavior following adolescent morphine exposure, such as increased inflammatory pain [63,64]. In these studies, morphine treatment was initiated during a period commonly defined as early adolescence in rodents (PND 21-34), whereas mice in the current study began treatment during mid-adolescence, which is defined as PND 34-46 [65–67]. Neurobehavioral sensitivity to psychoactive drugs has been shown to differ between stages of adolescence [66,68], thus, earlier developmental periods may be more susceptible to morphine-mediated effects on the immune system. Additionally, since females tend to enter puberty earlier than males [69–72], future experiments should consider sex-specific critical periods within adolescence.

A limitation of the current study is that only one dose of LPS was investigated. LPS dose-dependently induces the production of pro-inflammatory cytokines [28,30,73]. It is possible that the dose of LPS used in the current study was sufficient to induce some sickness behaviors without promoting a full immune response. Future studies could examine higher doses of LPS to investigate the impact of adolescent morphine exposure in more severe inflammatory states, like sepsis. Implementing high-speed videography and machine learning approaches in future research would facilitate objective measurement of physical discomfort in rodents, such as piloerection, ptosis, ear changes, orbital tightening, and nose/cheek flattening [74,75]. Lastly, the scope of the current study was limited to behavioral and physiological measurements; thus, it is unclear whether adolescent morphine exposure altered neuroimmune response to LPS on a molecular level. Future research may seek to quantify microglial activation, the presence of peripheral macrophages in the brain, and the expression of pro- and anti-inflammatory cytokines to assess short-term and long-term neuroinflammatory states following LPS exposure.

Further investigation is needed to fully elucidate the long-term neuroimmune effects of early-life opioid exposure. Greater insight into how opioids affect immune system signaling during sensitive developmental periods is important for enhancing health outcomes in vulnerable populations impacted by the opioid crisis.

